# Reprogramming of breast tumor-associated macrophages with modulation of arginine metabolism

**DOI:** 10.1101/2023.08.22.554238

**Authors:** Veani Fernando, Xunzhen Zheng, Vandana Sharma, Saori Furuta

## Abstract

HER2+ breast tumors have abundant immune-suppressive cells, including M2-type tumor associated macrophages (TAMs). While TAMs consist of the immune-stimulatory M1-type and immune-suppressive M2-type, M1/M2-TAM ratio is reduced in immune-suppressive tumors, contributing to their immunotherapy refractoriness. M1 vs. M2-TAM formation depends on differential arginine metabolism, where M1-TAMs convert arginine to nitric oxide (NO) and M2- TAMs convert arginine to polyamines (PAs). We hypothesize that such distinct arginine metabolism in M1- vs M2-TAMs is attributed to different availability of BH_4_ (NO synthase cofactor) and that its replenishment would reprogram M2-TAMs to M1-TAMs. Recently, we reported that sepiapterin (SEP), the endogenous BH_4_ precursor, elevates the expression of M1- TAM markers within HER2+ tumors. Here, we show that SEP restores BH_4_ levels in M2-TAMs, which then redirects arginine metabolism to NO synthesis and converts M2-TAMs to M1-TAMs. The reprogrammed TAMs exhibit full-fledged capabilities of antigen presentation and induction of effector T cells to trigger immunogenic cell death of HER2+ cancer cells. This study substantiates the utility of SEP in metabolic shift of HER2+ breast tumor microenvironment as a novel immunotherapeutic strategy.

## Introduction

Tumor associated macrophages (TAMs) are a heterogenous group of macrophages within solid tumors, comprising up to 50% of the cell mass of tumors (Obeid et al., 2013). They largely account for the immunosuppressive nature of the tumor microenvironment (TME) owing to the predominance of the immune-suppressive M2 type over the immune-stimulatory M1 type (Ayoub et al., 2019; Chanmee et al., 2014). M1-TAMs are classically activated macrophages that induce pro-immunogenic anti-tumor responses within the TME. In response to pro-inflammatory stimuli, such as LPS, IFNγ, IL12 and GM-CSF, nascent (M0) macrophages are polarized to M1-TAMs and induce immune-stimulatory Th1 responses via antigen presentation and secretion of immunogenic chemokines (CXCL9, CXCL10) (van Dalen et al., 2018; Wang et al., 2014). M1- TAMs also produce high levels of tumoricidal nitric oxide (NO) and reactive oxygen species (ROS) as well as pro-inflammatory cytokines (IL12, IL6,IL1β, TNFα) (Heusinkveld and van der Burg, 2011; Martinez et al., 2006; Raggi et al., 2017). On the contrary, M2-TAMs are alternatively activated macrophages that exert anti-inflammatory pro-tumor effects. Type 2 immunogenic stimuli, such as IL4, IL13 and M-CSF, trigger M2-TAM polarization, leading to the induction of immune-suppressive Th2 responses. M2-TAMs also produce a large amount of polyamines (PAs), polycations that promote secretion of anti-inflammatory cytokines, namely, IL10 and TGFβ, to facilitate tumor growth (Chanput et al., 2013; Lolo et al., 2017; Martinez and Gordon, 2014; Zhang et al., 2016).

Reduction of M1/M2-TAM ratio, as seen in immune-suppressive tumors such as human epidermal growth factor receptor 2 (HER2)-positive breast tumors, aggravates tumor growth and therapy resistance. Conversely, increased M1/M2-TAM ratio improves tumor prognosis and therapeutic response (Boutilier and Elsawa, 2021; Duan and Luo, 2021; Honkanen et al., 2019). Thus, reprogramming M2-TAMs to M1-TAMs is an emerging therapeutic strategy currently at investigational stages. Nevertheless, most of such endeavors utilize pro-inflammatory agents (LPS, IFNγ, TNFα, CD40 agonists, and IL12) that could cause systemic toxicity *in vivo* and fail in clinical utilization (Cai et al., 2021; Eisel et al., 2019). Thus, it is essential to develop a therapeutic strategy to reprogram TAMs with little side effects on patients. Recently, metabolic modulation has emerged as a safe approach to reprogram TAMs, based on the finding that different TAM subtypes exhibit distinct metabolic profiles (Fiaschi and Chiarugi, 2012; Liu et al., 2021). One such amenable metabolism is arginine catabolism (Massi et al., 2007). We previously reported that arginine catabolism in the breast was shifted from the immune-stimulatory NO synthesis pathway towards the immune-suppressive PA synthesis pathway during breast tumor progression, especially for HER2-positive tumors. This was largely attributed to NO synthase (NOS) dysfunction in the TME due to oxidative degradation of the essential enzyme cofactor, tetrahydrobiopterin (BH_4_). Thus, replenishment of BH_4_ in tumors by supplementing the endogenous precursor sepiapterin (SEP) effectively redirected arginine metabolism from PA to NO synthesis, reprogramed M2 TAMs to M1 TAMs, and suppressed tumor cell growth (Ren et al., 2019; Zheng et al., 2020).

In the present study, we explored the mechanisms by which modulation of arginine metabolism could reprogram TAMs and determined the therapeutic potentials of SEP for HER2- positive breast cancer. We found that bimodal arginine metabolic pathways leading to NO vs. PA synthesis are not only the consequences, but also the drivers of M1 vs. M2 macrophage polarization. Activation of enzymatic pathways for NO or PA synthesis as well as these metabolites were essential to polarize nascent M0 macrophages to M1 vs. M2 types, respectively. We further demonstrated that SEP-treated M2-TAMs not only elevated the expression of M1 TAM markers, but also exhibited full-fledged M1 TAM functionalities, including the elevated capacities of antigen presentation and cytotoxic T cell activation. When co-cultured with these M1 reprogrammed TAMs as well as activated T cells, cancer cells underwent immunogenic cell death (ICD) indicated by the production of damage-associated molecular patterns (DAMPs) and apoptotic markers. Such strong pro-immunogenic, anti-tumor effects of SEP were verified using animal models of spontaneous HER2-positive mammary tumors. These results strongly suggest the immunotherapeutic potentials of SEP-mediated metabolic shift of TAMs for HER2-positive breast cancer.

## Results

### M1 and M2 TAMs are distinguished by the preferential production of NO vs. PAs through differential arginine metabolism

Circulating monocytes that have entered the TME differentiate into nascent (M0) TAMs and then polarize into different subtypes based on the environmental cues (de Sousa et al., 2019; Heusinkveld and van der Burg, 2011; Juhas et al., 2015; Kalish et al., 2017; Malyshev and Malyshev, 2015). M1 and M2-TAMs are the two major subsets that represent the opposing ends of a spectrum in terms of morphology, metabolism and functions (Italiani and Boraschi, 2014; Martinez and Gordon, 2014; Tarique et al., 2015; Wang et al., 2014). We utilized *in vitro* TAM models that represent M0, M1 and M2 subtypes derived from THP–1 monocytic cells **(Figure 1A)**(Li et al., 2016) as well as peripheral blood mononuclear cells (PBMCs) **(Figure 1B)**. To distinguish between different TAM subsets, we profiled different TAMs based on the morphologies, metabolisms, and functional signatures. Phalloidin staining of filamentous actin as well as scanning electron microscopy (SEM) imaging of TAMs showed that M0-type appeared as sparsely clustered spherical cells, whereas M1-type manifested as more elongated, spindle-shaped cells. M2-type, on the other hand, formed highly clustered and more spread morphology as reported previously **(Figure 1C)**(Bertani et al., 2017; Lendeckel et al., 2022). M1- and M2-TAMs were more quantitively distinguished by their unique marker expression (Heinrich et al., 2017; Maess et al., 2010; Martinez et al., 2006). M1-TAMs showed significantly higher levels of TNFα and TLR2 and little expression of CD206. In contrast, M2-TAMs showed little or low expression of TNFα and TLR2, but higher levels of CD206. Interestingly, M0-TAMs expressed both M1 and M2 markers, although at lower levels, attesting to their bipotency prior to polarization (**Figures 1D-F**). Furthermore, M1- vs. M2-TAMs are characterized by their differential arginine catabolism (Massi et al., 2007). M1-TAMs metabolize arginine via NOS2 to produce NO for anti–tumor activities. Conversely, M2 TAMs metabolize arginine by arginase 1 (Arg1) and then by ornithine decarboxylase 1 (ODC1) to produce PAs for pro–tumor activities (Boucher et al., 1994; Geelhaar- Karsch et al., 2013; Rodriguez et al., 2017; Zheng et al., 2020). Consistently, we observed that M1-TAMs preferentially produced NO over PAs while M2-TAMs preferred PAs over NO, as indicated by the differential NO/PA ratios **(Fig 1G)**.

**Figure 1:**
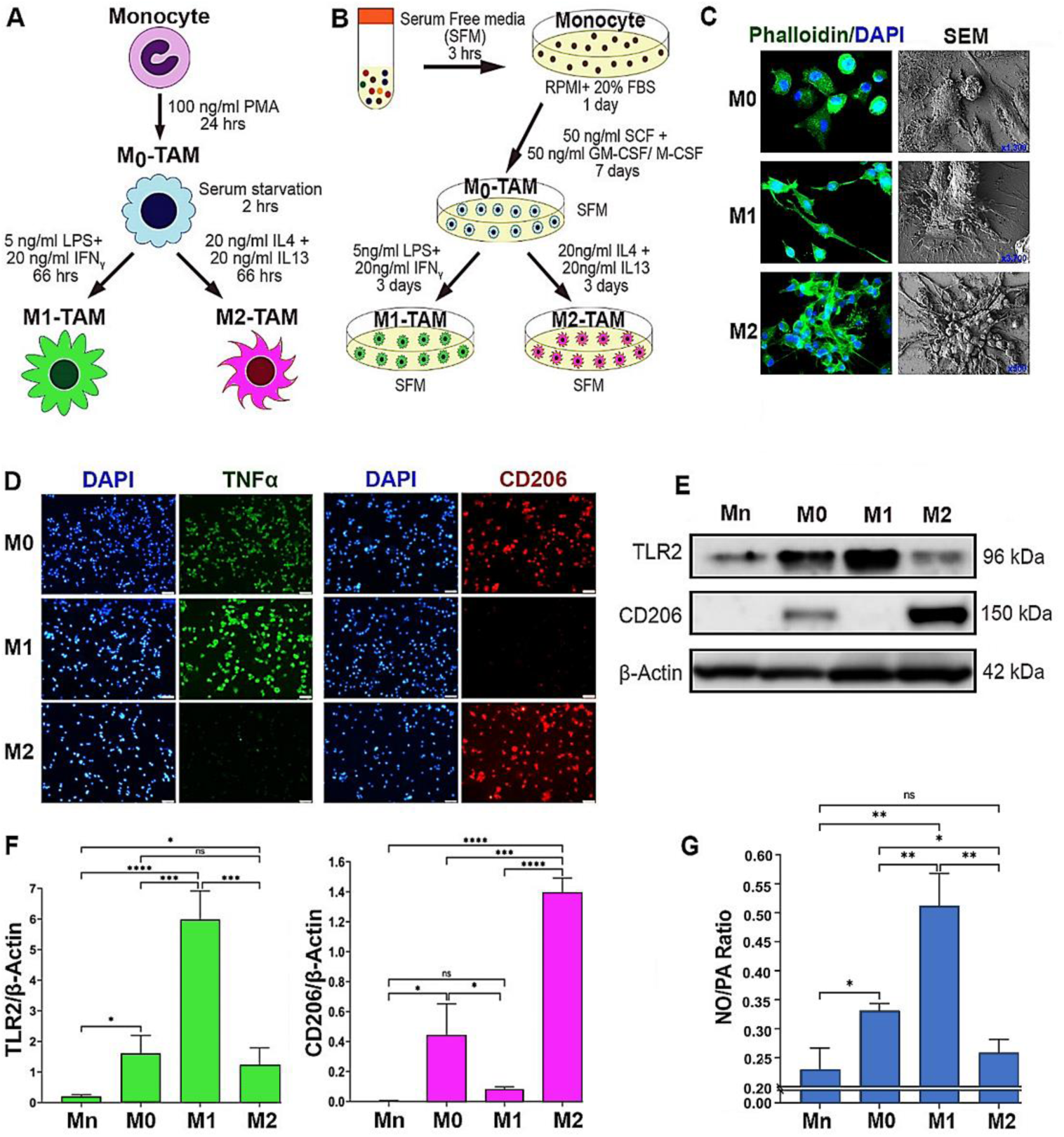
M1 vs. M2 TAMs are distinguished by the preferential production of NO vs. PAs through differential arginine metabolism. **A, B**) Schematic representation of the polarization protocols of TAMs derived from THP–1 human monocytic cell line (A) & PBMC (B). THP–1 monocyte (Mn) was treated with phorbol myristate aetate (PMA) for differentiation to inactive macrophages (M_0_). PBMC-derived Mn was treated with SCF+GM-CSF (leading to M1) or M-CSF (leading to M2) for differentiation to M_0_. For M1 polarization, M_0_–TAMs were treated with LPS and IFNγ; for M2 polarization, M_0_–TAMs were treated with IL4 plus IL13. **C**) Representative images of phalloidin staining (left, n=3) & scanning electron microscopy (SEM, right, n=3) imaging of M0, M1 and M2-TAMs. Green: phalloidin; blue: DAPI. SEM images are shown at different magnifications; M0: x1300), M1: x3700), and M2: x500. **D**) Immunofluorescence images of TAM subsets (n=3) stained for M1 (green, TNFα) vs. M2 markers (red, CD206) and counterstained with DAPI (blue). Scale bars: 50μm. **E**) Western blot analysis (n=3) of THP–1 derived Mn, M0, M1 and M2-TAMs for the expression of TLR2 (M1 marker) vs. CD206 (M2 marker). β–Actin was used as the internal loading control. **F**) Quantification of the western results of the expression of TLR2 (left) and CD206 (right) normalized against β–Actin and presented as fold differences. **G**) NO to PA ratios in THP–1 derived TAM subsets (n = 6) measured with ELISA. Error bars: ±standard errors of mean (SEM). *, p ≤ 0.05; **, p ≤ 0.01; ***, p ≤ 0.001; ****, p ≤ 0.0001 and ns, p >0.05.

### SEP reprograms M2-TAMs to M1-TAMs by downmodulating PAs but upregulating NO synthesis

We previously reported that administration of SEP, the endogenous precursor of the NOS cofactor BH_4_, could redirect arginine metabolism from PA to NO synthesis in the breast TME, which in turn upregulated M1-TAM marker expression while downmodulating M2 TAM markers (Zheng et al., 2020). In the present study, we sought to investigate whether M2 TAMs treated with SEP would indeed acquire M1-TAM functionalities using macrophages derived from THP–1 cells. We polarized these macrophages to M1 vs. M2 types and treated them with SEP (100 μM), in comparison to the equal volume of DMSO as vehicle control, for 3 days and determined their phenotypic profiles. M1-TAMs treated with SEP (100 μM) and M2-TAMs treated with LPS (5 ng/ml) plus IFNγ (20 ng/ml) were used as positive controls.

First, to determine the bioactivity of SEP, we measured BH_4_ production. Although the endogenous BH_4_ levels were significantly lower in M2-TAMs than M1-TAMs, SEP treatment elevated BH_4_ in both M1 and M2-TAMs to almost equal levels (**Figure 2A**). SEP-treated M2- TAMs showed M1-TAM-like morphology characterized by isolated spindle-shapes **(Fig 2B).** Consistent with our previous report (Zheng et al., 2020), SEP treated M2-TAMs showed large increases in M1-TAM markers (TLR2, TNFα, IL1β, and IL6) and decreases in M2 TAM markers (CD206 and TGF β), although to a lesser degree than positive control (LPS plus IFNγ) **(Fig 2C, F)**. We then tested the effects of SEP on arginine metabolism of TAMs. Consistent with our previous findings (Zheng et al., 2020), M1-TAMs produced 3 fold higher levels of NO than M2- TAMs, while M2-TAMs produced 4 fold higher levels of PAs than M1-TAMs. Thus, NO/PA ratio of M1-TAMs was over 8 fold higher than that of M2-TAMs. Upon SEP treatment, however, NO/PA ratio in M2-TAMs increased by 3 fold, while it did not change in M1-TAMs **(Fig 3A)**. These results suggested that SEP redirected the arginine metabolism of M2-TAMs from PA to NO synthesis.

**Figure 2:**
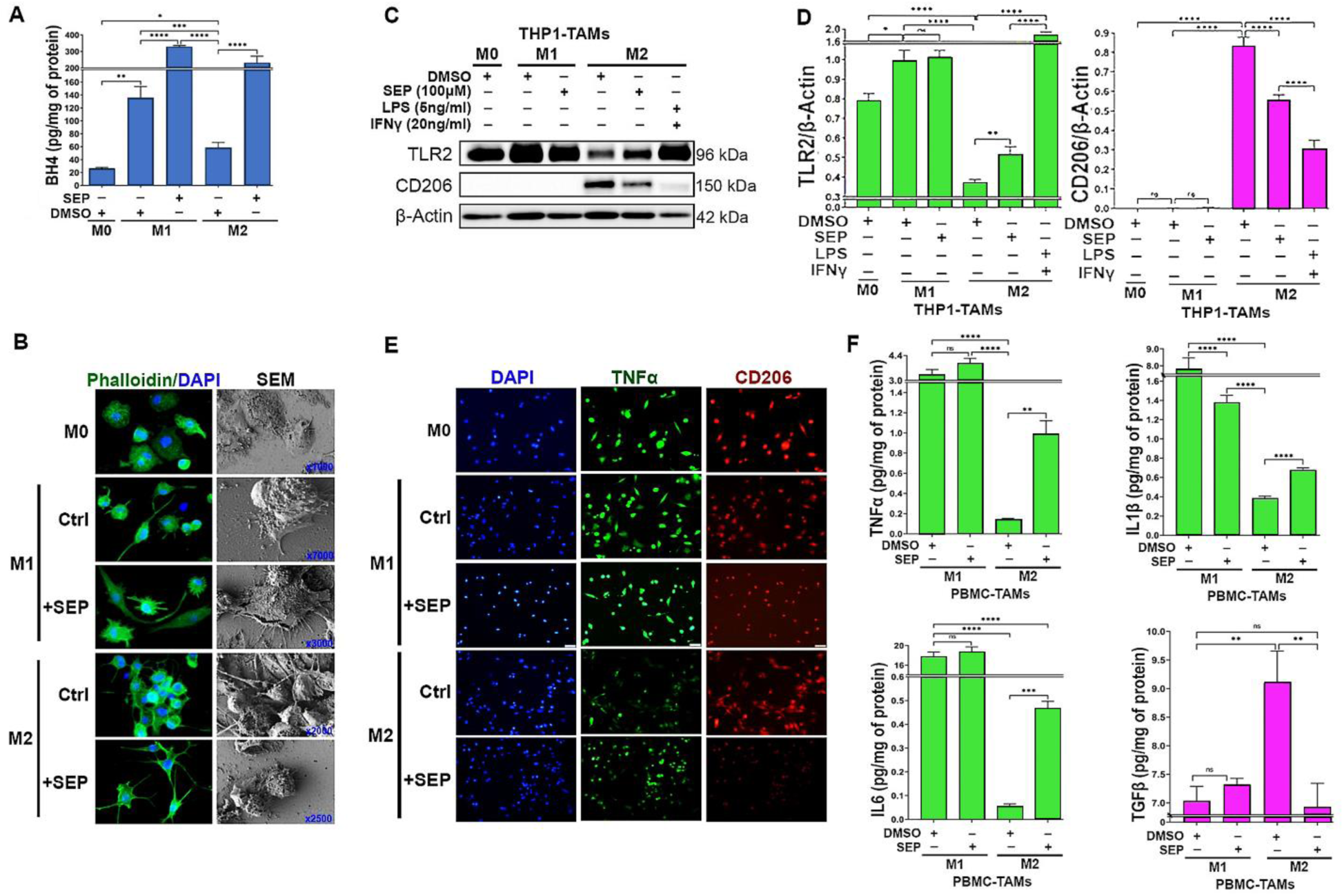
SEP elevates M1 marker expression in M2 TAMs. **A**) Levels of BH_4_ produced by THP–1 derived M0, M1, and M2-TAMs after treated with vehicle (DMSO) or SEP (100μM) for 3 days (n=6). BH_4_ levels were measured with ELISA and normalized against the total protein levels. One way ANOVA with post hoc test (Tukey test) was performed to measure the significance of mean difference between treatment groups. **B**) Phalloidin staining and SEM imaging of THP–1 derived M0, M1 and M2-TAMs treated as in **A**) (n=3). SEM images are shown at different magnifications. **C**) Western blot analysis of TAM subsets treated with vehicle or SEP as in **A**), and β–Actin was used as the internal loading control (n=5). **D**) Quantification of the western results based on the expression of TLR2 (M1 marker) vs. CD206 (M2 marker) normalized against β–Actin signal and presented as fold differences. **E**) Immunofluorescence imaging of THP–1 derived TAMs after treatments as shown above and stained for M1 marker (green, TNFα) vs. M2 marker (red, CD206) and counterstained with DAPI (blue)(n=3). **F**) The levels of secreted cytokines, Type 1: TNFα (top left), IL1β (top right), and IL6 (bottom left) vs. Type 2: TGFβ (bottom right), for M1 vs. M2 TAMs treated with vehicle vs. SEP (n=6) measured with ELISA. Error bars: ±SEM. *, p ≤ 0.05; **, p ≤ 0.01; ***, p ≤ 0.001; ****, p ≤ 0.0001 and ns, p >0.05.

**Figure 3:**
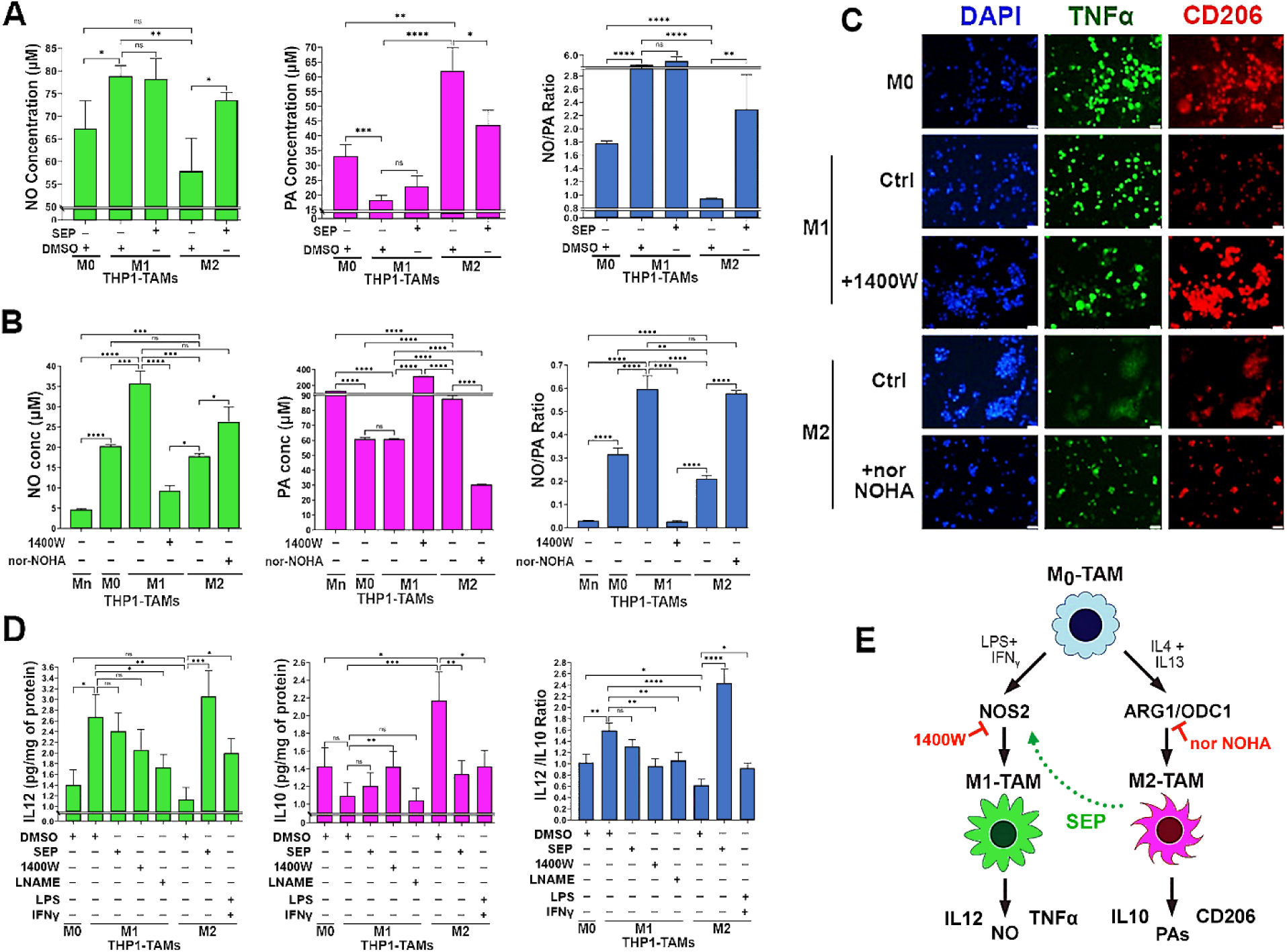
SEP redirects arginine metabolism from PA to NO synthesis in M2 TAMs, while rendering them M1-TAM phenotype. **A**) Levels of NO (left), PAs (middle) and NO/PA ratios for THP–1 derived M0, M1 and M2- TAMs after treated with DMSO (vehicle) and SEP (100 μM) for 3 days (n = 5). B) One way ANOVA with post hoc Tukey test was used for statistical analysis. Error bars: ±SEM. GraphPad Prism Version 9.5.1. was used to perform all statistical analyses. **B**) Levels of NO (left), PAs (middle) and NO/PA ratios for THP–1 derived Mn, M0, M1 and M2-TAMs after treated with DMSO (vehicle), NOS2 inhibitor, 1400W (50 μM), or Arginase 1 (Ang1) inhibitor, nor NOHA (50 μM) for 3 days (n = 5). Note the significant decrease of NO level in 1400W-treated M1-TAMs and significant decrease of PA level in nor-NOHA-treated M2-TAMs. **C**) Immune fluorescence imaging of THP–1 derived M0, M1 and M2-TAMs stained for M1 marker (green, TNFα) vs. M2 marker (red, CD206) and counterstained with DAPI (blue). M1-TAMs were treated with DMSO (Control: Ctrl) or NOS2 inhibitor (100μM 1400W), whereas M2-TAMs were treated with DMSO (Ctrl) or ARG1 inhibitor (50μM nor-NOHA) for 3 days (n=3). **D**) Levels of Type 1 cytokine IL12 (left) and Type 2 cytokine IL10 (middle) as well as IL12/IL10 ratios for THP-1 derived TAM subsets measured with ELISA. M1-TAMs were treated with DMSO or NOS inhibitors, 1400W (50 μM) and LNAME (2.5 mM). M2-TAMs were treated with DMSO, SEP (100 μM) or positive control LPS (5ng/ml) plus IFNγ (20ng/ml) for 3 days (n=6). The cytokine levels were measured using ELISA and normalized against the total protein levels. Error bars: ±SEM. *, p ≤ 0.05; **, p ≤ 0.01; ***, p ≤ 0.001; ****, p ≤ 0.0001 and ns, p >0.05. **E**) Working scheme for the induction of M1 vs. M2 polarization by activation of NOS2 vs. ARG1/OCD1 pathways and M2-to-M1 reprogramming by SEP.

To explore how SEP-mediated modulation of arginine metabolism had influenced TAM polarization, we determined the relevance of NO vs. PA synthesis pathways to M1- vs. M2-TAM polarization, respectively. We inhibited NO synthesis pathway with a specific NOS2 inhibitor 1400W (50 μM), while inhibiting PA synthesis pathway with a specific ARG1 inhibitor nor-NOHA (50 μM). As expected, 1400W significantly downmodulated NO levels in M1-TAMs, whereas nor-NOHA downmodulated PA levels in M2-TAMs (**Fig. 3B**). Concomitantly, 1400W treated M1-TAMs showed decrease of M1-TAM markers, TNFα and IL12, but increase of M2- TAM markers, CD206 and IL10. In contrast, nor-NOHA-treated M2-TAMs showed decreases of M2-TAM markers, but increases of M1-TAM markers (**Fig. 3C, D**). These results suggest that NOS2 vs. ARG1 functions are essential for the polarization to M1- vs. M2-TAMs, respectively (**Fig. 3E**). We further tested the relevance of NO vs. PA per se to M1- vs. M2-TAM polarization (Zheng et al., 2020), respectively. While we and others observed that NO and PAs were produced as the consequence of M1 vs. M2 polarization (Massi et al., 2007), we suspected that both metabolites could also be the ‘driver’ of M1 vs. M2 polarization. First, we treated nascent M0- TAMs with SEP. NO donors (GSNO and SNAP) or PA (Spermine) and measured M1 and M2 marker epression. M0-TAMs treated with SEP or NO donors significantly elevated M1 markers (TLR2, STAT1, pSTAT1_S727_, and IL12), while decreasing M2 marker (CD163, STAT3, and IL10) **(Fig 4 A-C, Appendix Figure S1)**. Next, we treated M1-TAMs with a NO scaventer cPTIO and M2-TAMs with an inhibitory PA analogue DENSPM. We showed that cPTIO effectively inhibited M1 markers, while DENSPM inhibited M2 markers (**Fig. 4 D-F, Appendix Figure S2**). These results suggest that NO and PAs are indeed drivers of M1- vs. M2-TAM polarization, accounting for the mechanism of SEP-induced reprogramming of M2- to M1-TAMs.

**Figure 4:**
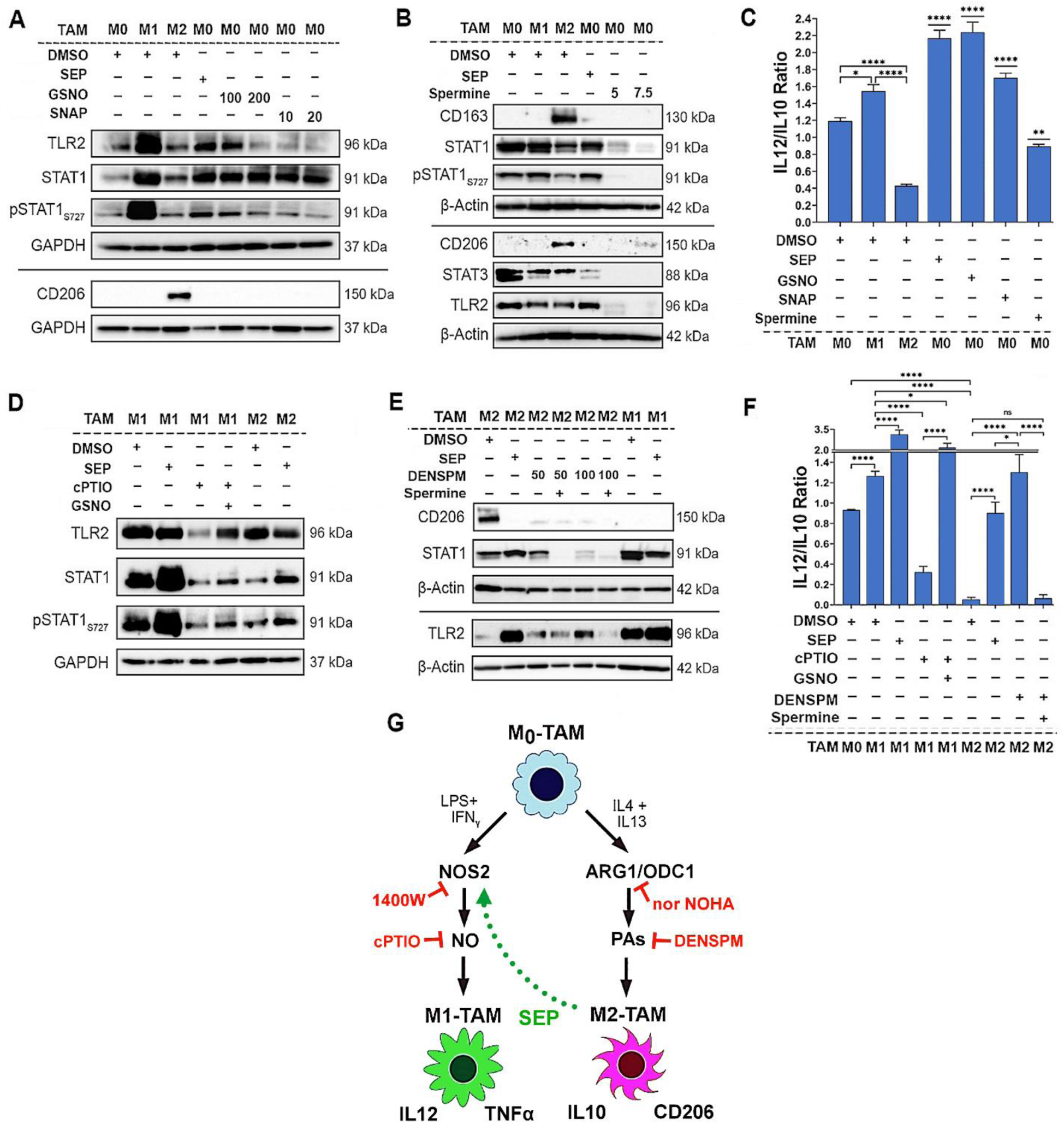
Arginine metabolites, nitric Oxide and polyamines, drive M1 and M2-TAM polarization, respectively. **A**) Western blot analysis on M1 markers: TLR2, STAT1 and pSTAT1_S727_ vs. M2 marker: CD206 in THP1–derived M0-TAMs treated with DMSO (vehicle control), SEP (100 μM), or NO donor (GSNO [100 μM, 200 μM] and SNAP [10 μM, 20 μM]) in comparison to M1- and M2-TAMs (n=4). GAPDH was used as the internal loading control. **B**) Western blot analysis on M1 markers: STAT1, and TLR2, vs. M2 marker: CD206 in M0-TAMs treated with DMSO (vehicle control), SEP (100 μM), or PAs (5 or 7.5 mM Spermine) in comparison to M1- and M2-TAMs (n=4). β– Actin was used as the internal loading control. (For quantification of **A** and **B**, see **Appendix Figures S1**.) **C**) The ratios of IL12/IL10 secreted by THP–1 derived M0-TAMs treated with DMSO, SEP, NO donors and PAs in comparison to M1- and M2-TAMs. **D**) Western blot analysis on M1 markers: TLR2, STAT1 and pSTAT1_S727_ in M1-TAMs treated with NO scavenger (50 μM cPTIO) with and without NO donor (100 μM GSNO) and M2-TAMs treated with DMSO or SEP (n=4). GAPDH was used as the internal loading control. **E**) Western blot analysis on M1 markers: TLR2, STAT1 and pSTAT1_S727_ vs. M2 marker: CD206 in THP1-derived M2-TAMs treated with PA analog (50 μM, 100 μM DENSPM) and PAs (5 mM Spermine) and M1-TAMs treated with DMSO or SEP (n=4). β–Actin was used as the internal loading control. (For quantification of **D** and **E** see **Appendix Figures S2**.) **F**) The ratios of IL12/IL10 secreted by THP–1 derived M1- and M2-TAMs with treatment combinations shown in **D**) and **E**). Error bars represent ± SEM. GraphPad Prism Version 9.5.1. was used to perform all statistical analyses. *, p ≤ 0.05; **, p ≤ 0.01; ***, p ≤ 0.001; ****, p ≤ 0.0001 and ns, p >0.05. **G**) Scheme for the induction of M1 vs. M2 polarization by NO vs. PAs and M2-to-M1 reprogramming by SEP.

### SEP-treated M2 TAMs exhibit enhanced antigen presentation capabilities

To validate these reprogrammed TAMs are indeed functional, we measured their antigen presentation capabilities. Macrophages are professional antigen presenting cells (APC) that activate the adaptive immune system upon detection of foreign antigens (Barker et al., 2002; Martín-Orozco et al., 2001; Muntjewerff et al., 2020). M1-TAMs could present neoantigens derived from tumor cells towards CD4+Type I T helper cells (Th1) via HLA-DR (MHCII) receptors, which in turn activates CD8+ cytotoxic T cells (Mills and Ley, 2014; Pan et al., 2020; Wang et al., 2019). M2-TAMs, on the other hand, suppress cytotoxic T cell activity by expressing the tolerogenic HLA-G antigen (Boutilier and Elsawa, 2021; Contini et al., 2020). To measure antigen presentation capability, we used OVA _323-339_ (OVA) peptide, an antigenic epitope of HLA-DR (McFarland et al., 1999; Westerberg et al., 2003) that would be presented via the cell surface HLA-DR on M1-TAMs towards the T cell receptor (TCR) on Th1 cells **(Fig 5A)**. We then measured the levels of the cell surface HLA-DR which would emerge after antigen binding. As expected, in the absence of OVA, only basal level of surface HLA-DR was detected for all the TAM subsets. Upon being pulsed with OVA, however, M1-TAMs expressed significantly higher levels of the cell surface HLA-DR than M2-TAMs. Conversely, SEP treatment dramatically elevated the cell surface HLA-DR levels in M2-TAMs (5 fold) as well as M1-TAMs (1.5 fold). Overall, these findings demonstrate that SEP significantly elevated the antigen presentation capability of M2-TAMs to the levels equivalent to that of M1-TAMs (**Fig. 5B-D**).

**Figure 5:**
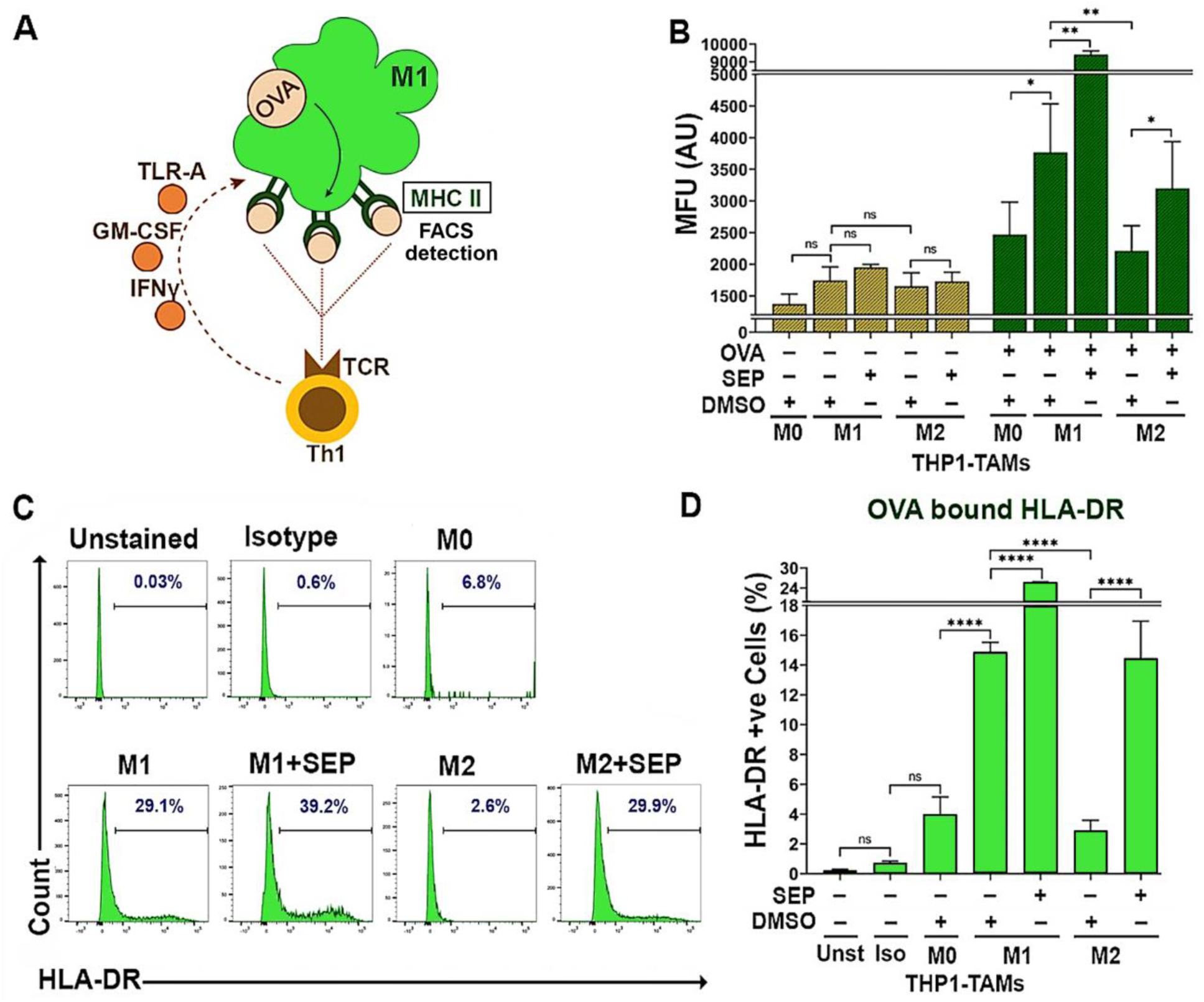
M2-TAMs treated with SEP show increased antigen presentation activities. **A**) The scheme of measuring antigen presentation activities of TAMs. Once M1 macrophage is pulsed with OVA_323-339_ peptide, it phagocytoses and presents the epitope through the cell surface MHC II. The level of cell surface MHC II, representing antigen presentation activity, is detected by FACS. (The presented epitope is then recognized by T cell receptor (TCR) on Th1 T cells to trigger immunogenic responses.) **B**) Mean fluorescence intensity (MFU) of cell surface HLA-DR (MHC II) after pulsed with or without OVA_323-339_ peptide (20μg/ml for 2 hours)) on THP–1 derived M_0_, M1 and M2-TAMs pre-treated with DMSO or 100 μM SEP. Two sample t-tests (unpaired) were performed for pairwise comparison. **C, D**) Percentages of TAM subsets treated as in **B**) that expressed cell surface HLA-DR (bound by OVA peptide) presented as histograms (**C**) and quantification (**D**). Unstained (Unst) and isotype (Iso) controls were used (n=5). Error bars: ± SEM. *, p ≤ 0.05; **, p ≤ 0.01; ***, p ≤ 0.001; ****, p ≤ 0.0001 and ns, p >0.05.

### SEP-treated M2-TAMs effectively activate cytotoxic T cells

We next tested whether these reprogramed TAMs could indeed activate cytotoxic T cells. Cytotoxic T cells are the major component of the adaptive immune system and the executors of anti-tumor immune responses. Cytotoxic T cells are activated through antigen presentation and induce cancer cell death by releasing cytotoxic proteins such as granzymes, perforin and IFNγ (Golstein and Griffiths, 2018; Jorgovanovic et al., 2020; Weigelin et al., 2021; Weigelin and Friedl, 2022). M1-TAMs are able to activate cytotoxic T cells, while M2-TAMs instead inhibit their activation (Mills and Ley, 2014). To determine T cell activation, we measured their IFNγ production, cell proliferation, and CD107a surface expression (degranulation) (Cheng et al., 2023; Cohnen et al., 2013; Lorenzo-Herrero et al., 2019; Mayer et al., 2019) **(Fig 6A)**. To this end, we devised two different co-culture systems: indirect (transwell-based) and direct methods, to co-culture TAMs, T cells and breast cancer cells. The indirect method was used to co-culture THP- 1-derived TAMs, PBMC-derived T cells and cancer cells, whereas direct method was used to co-culture PBMC-derived TAMs and T cells along with cancer cells **(Fig 6B)**. Here, we specifically focused on HER2+ breast cancer cells based on our previous studies demonstrating the pathogenesis of this cancer subtype responsive to dysregulated NO levels and their great sensitivity to SEP treatment (Ren et al., 2019; Ren et al., 2021; Zheng et al., 2020). We observed that in these triple co-cultures, M1-TAMs, but not M2-TAMs, strongly induced T cells to produce IFNγ, proliferate, and express cell surface CD107a **(Fig 6C-G)**. However, SEP-treated M2-TAMs dramatically elevated these three activation markers in co-cultured T cells. SEP-treated M1- TAMs, however, did not exhibit further increase in T cell activation, indicating the existence of certain threshold levels. These results strongly suggest that M2-TAMs treated with SEP could strongly activate cytotoxic T cells, as the result of their functional reprogramming to M1-TAMs.

**Figure 6:**
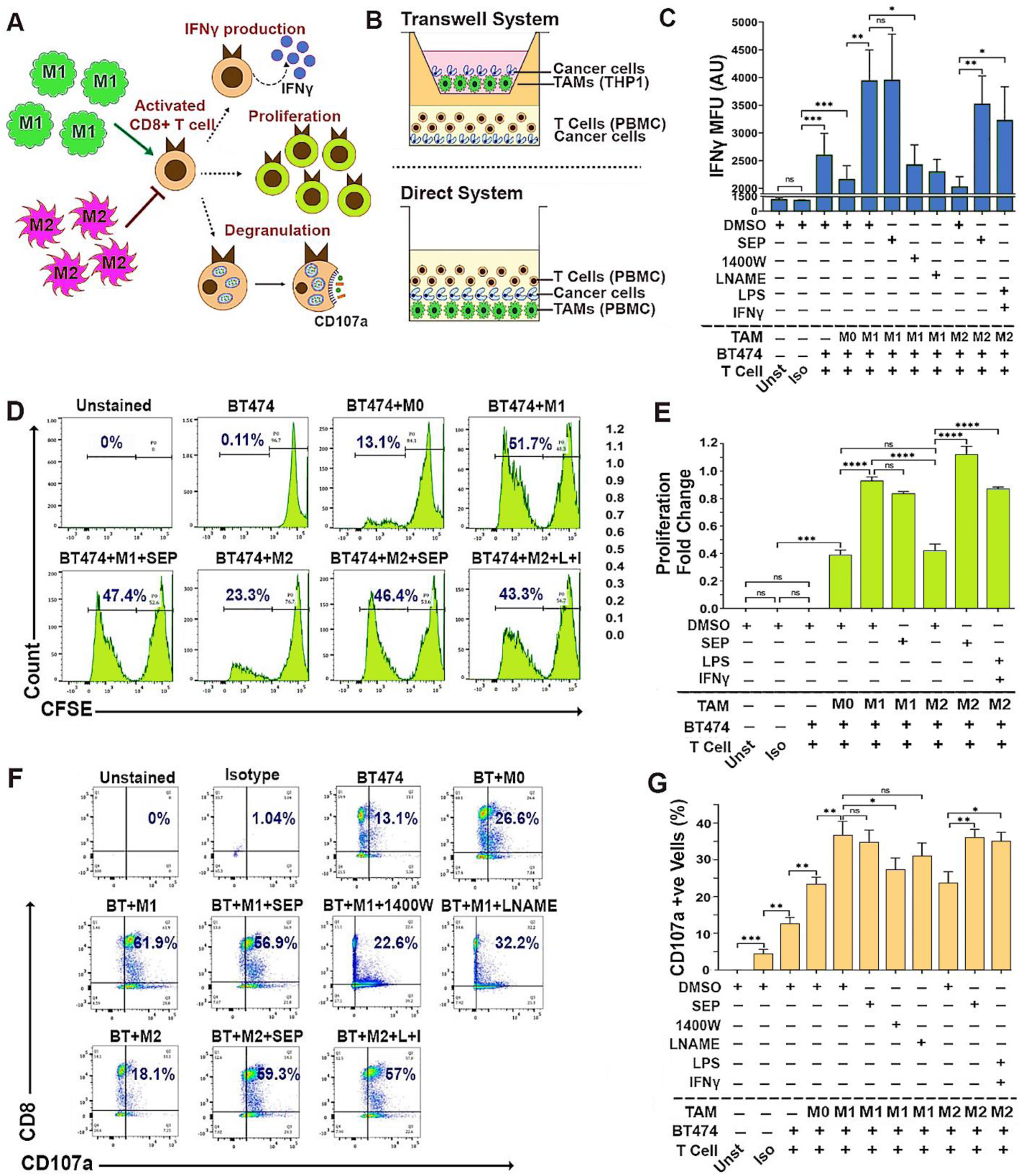
SEP treated M2-TAMs activate cytotoxic T cells. **A)** Scheme of detection methods for activation of cytotoxic CD8+ T cells, based on IFNγ production, proliferation, and degranulation indicated by cell surface CD107a expression. **B**) Schemes of the TAM-T cell-cancer cell co-cultures (2:2:1 ratio) using Transwell system (top) and direct system (bottom). (Top) THP–1 derived TAMs, PBMC derived T cells, and BT474 breast cancer cells co-cultured using Transwell. (Bottom) PBMC derived autologous TAMs and T cells were directly co-cultured with BT474 cells. **C**) FACS-detected IFNγ expression levels in CD8+ T cells after co-cultured with BT474 cells and PBMC derived TAM subsets pretreated with DMSO or 100 μM SEP (n=6). Positive control: M2-TAMs treated with LPS and IFNγ. Negative control: M1-TAMs treated with 1400W and LNAME. T cells were gated for CD3 and CD8 expression. IFNγ levels are shown as MFU. **D**) Detection of proliferation of cytotoxic T cells co-cultured as in **C**) based on the dilution of CFSE signals through cell doubling (n=6). Percentages of proliferating T cells (CFSE low) are highlighted in the histogram. **E**) Cytotoxic T cell proliferation shown as fold change of proliferating cells (CFSE low) with respect to non-proliferating cells (CFSE high). **F**) Cell surface CD107a (degranulation marker) expression on cytotoxic T cells directly co-cultured as in **B**). Percentages of CD107a+ CD8+ cells are shown in the plots. **G**) Quantification of the percentages of CD107a+ CD8+ cells in co-cultures. Error bars: ± SEM. *, p ≤ 0.05; **, p ≤ 0.01; ***, p ≤ 0.001; ****, p ≤ 0.0001 and ns, p >0.05.

### SEP-treated M2-TAMs induce ICD in HER2-positive breast cancer cells

Cytotoxic activities of T cells could be determined by measuring the death of target cells. To validate the cytotoxic activity of T cells co-cultured with TAMs, HER2+ breast cancer cells (BT474 & SKBR3) and the conditioned media (CM) from co-cultures were analyzed for cell death markers. Cell cycle profiling demonstrated that cancer cells co-cultured with SEP treated M2- TAMs along with T cells showed a great increase in the SubG1 (apoptotic) population compared to those co-cultured with control M2-TAMs **(Fig 7A, B, Appendix Figure S3A, B)**. This observation was further confirmed by the large increases in cancer cells positive for Annexin V (total apoptotic cells) at both early (PI low) and late (PI high) stages of apoptosis after being co-cultured with SEP-treated M2-TAMs and T cells (**Fig. 7C-E**).

**Figure 7:**
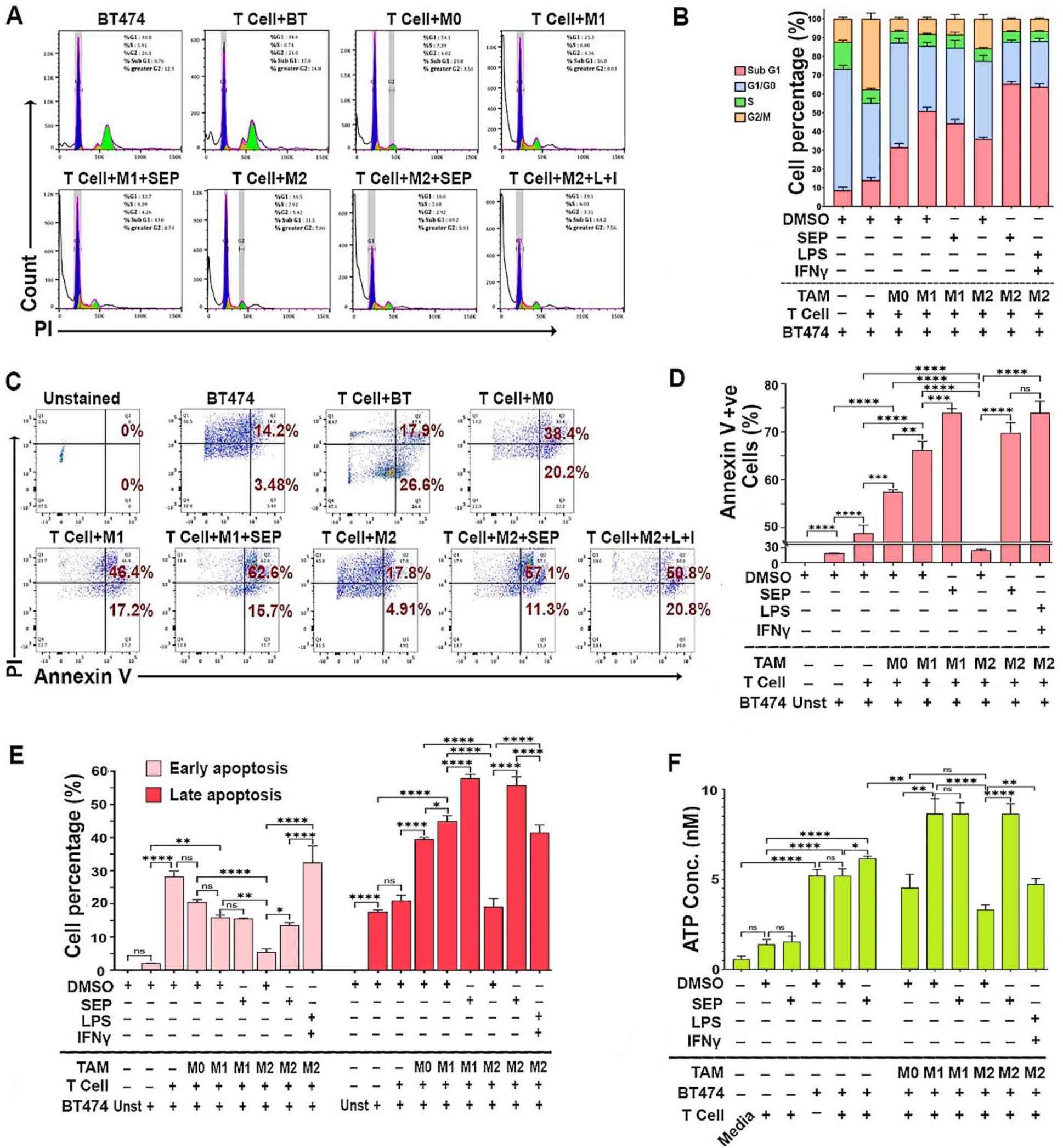
SEP treated M2-TAMs induce T cells to kill HER2+ breast cancer cells. **A**) Cell cycle analyses of BT474 cancer cells (CMFDA-labeled) co-cultured with PBMC derived TAM subsets, pretreated with DMSO (vehicle), SEP (100 μM), or LPS + IFNγ (positive control), along with T cells. Adherent cells (TAMs+BT474 cells) were dissociated, fixed in 70% ethanol for 3 hours, and stained with PI. BT474 cells were gated based on CMFDA signal and analyzed for the PI-stained DNA contents. **B**) Cell cycle distribution of BT474 cells co-cultured with TAMs and T cells as in **A**). Note the dramatic increase in SubG1 population in co-cultured with SEP treated M2-TAMs. (For quantification of Sub-G1 and G1/G0 populations, see **Appendix Figures S3A**.) **C**) Annexin V/PI staining of co-cultured BT474 cells to measure cell deaths (n=6). Viable cells: Annexin V-ve, PI-ve; early apoptotic cells: Annexin V +ve, PI-ve; late apoptotic cells: Annexin V +ve, PI +ve; and necrotic cells: Annexin V-ve, PI +ve. **D**) Percentage of total apoptotic (Annexin V +ve) cancer cells. **E**) Early and late apoptotic cancer cells. **F**) Levels of ATP secreted by BT474 cells in co-cultures. Secretion of ATP indicates immunogenic cell death of cancer cells. Note the dramatic increase in ATP secretion by cancer cells in co-culture with SEP-pretreated M2 TAMs. Error bars: ± SEM. *, p ≤ 0.05; **, p ≤ 0.01; ***, p ≤ 0.001; ****, p ≤ 0.0001 and ns, p >0.05.

To determine the mechanisms of cancer cell death after being co-cultured with SEP-treated TAMs along with T cells, we analyzed the CM for the levels of secreted ATP, a type of damage-associated molecular patterns (DAMPs) released from cells undergoing immunogenic cell death (ICD). DAMPs bind and activate the cognate cell surface receptors on phagocytic cells to mediate their own destruction (Jiang et al., 2022; Martins et al., 2014). CM of T cell only cultures contained the basal levels of ATP regardless of the treatment. The inclusion of cancer cells, however, increased the extracellular ATP levels by 2 folds, which were increased by SEP treatment by the additional 20%. The further addition of M1-TAMs, but not M0- or M2-TAMs, increased the secreted ATP levels by another 50%. Nevertheless, when M2-TAMs were pretreated with SEP, the addition of these M2-TAMs elevated the extracellular ATP to the levels equivalent to those of co-cultures with M1-TAMs. These findings demonstrated that SEP greatly enhanced cancer cell killing activities of adaptive immunity through ICD.

### Oral SEP treatment elevates M1-TAMs in TME and suppresses the growth of spontaneous MMTV-neu (HER2) mammary tumors

To validate the therapeutic efficacy of SEP, we gave SEP to MMTV-neu mice which were a mouse model of spontaneous HER2-positive mammary tumors. These animals developed single-focal tumors at the latencies of 6-14 months. Once tumors became palpable, animals were divided into the control (DMSO) vs. SEP (10 mg/kg) treatment groups and given the drug through ad libitum access to acidified drinking water for 6 weeks (**Figure 8A**). Tumor growth was measured twice a week using a caliper, and morbidity of animals were also observed. We saw about 50% reduction in the tumor growth curve as well as the sizes of the exercised tumors of SEP treated group with statistical significance (**Figures 8B, C**). We did not see any morbidity of animals due to treatments. To determine the immunogenicity of these tumors, we isolated TAMs and compared their M1- vs. M2-TAM marker expression. In SEP treated group, we saw large increases of M1 markers (CD80, IL12, and IFNγ) but decrease of a M2 marker, CD163. Another M2 marker, IL10, however, was not influenced by SEP treatment, indicating the persistence of a subtype of M2- TAMs (**Figures 8D, E**). Still, a large increase of M1-TAMs in SEP-treated tumors is expected to be the significant contributor to the tumor inhibitory effects of the drug. Such increase in M1-type macrophages and decrease in M2-type macrophages by SEP treatment were more consistent throughout different markers in tumors than spleens, indicating that the major targets of SEP were TAMs (**Figure 8E**). Our results altogether strongly suggest the potential of SEP as a novel immunotherapeutic agent for HER2-positive breast cancer.

**Figure 8:**
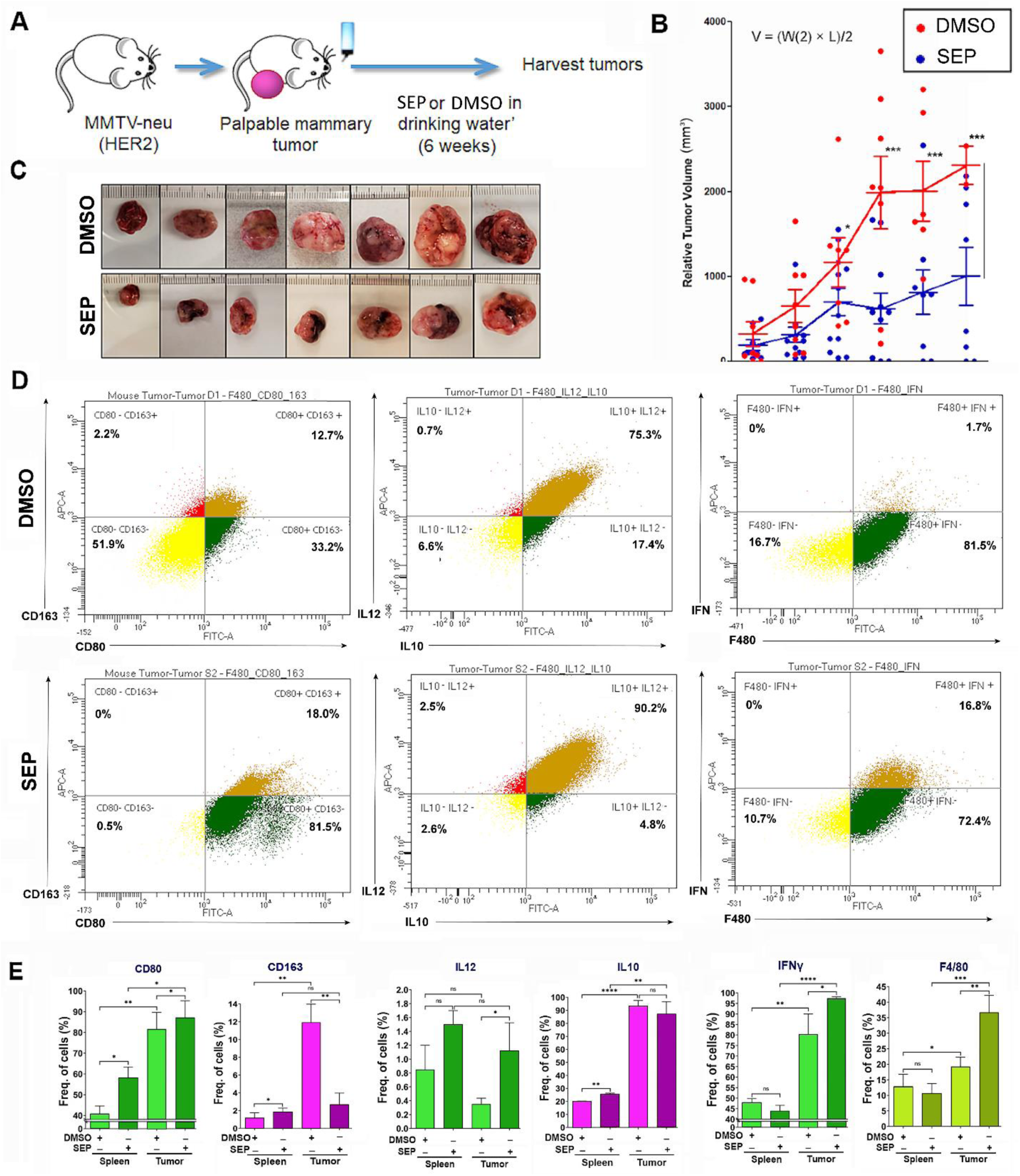
Oral SEP treatment promotes the immunogenicity of TAMs and suppresses the growth of spontaneous MMTV-neu (HER2) mammary tumors. **A**) Scheme of the experiment where MMTV-neu (unactivated) mice were allowed to develop palpable mammary tumors (tumor latency of 6-14 months) and given DMSO or SEP (10 mg/kg) in drinking water ad libitum for 6 weeks (n=7). **B**) Tumor growth was measured by caliper and the volume was determined (V = (W(2) x L)/2). **C**) Pictures of exercised tumors. **D**) Exercised tumors were analyzed for M1- vs. M2-TAM profiles (CD80 vs. CD163; IL12 vs. IL10; and IFNγ) by FACS. (Top row) DMSO-treated tumors; (bottom row) SEP-treated tumors. **E**) Quantification of the expression of M1- vs. M2-TAM markers as in **D**) in exercised tumors (n=6) in comparison to spleens (n=6) of the same animals.

## Discussion

TAMs are a group of heterogenous myeloid cells that manifest a great plasticity under the influence of varying signals from the TME. Although TAMs are broadly classified into the pro-inflammatory M1 vs. anti-inflammatory M2 types for simplicity’s sake, their given phenotypes could range within the spectrum. Both types of TAMs encompass distinct metabolic profiles depending on their specific needs for energy production. M1 TAMs tend to be localized in hypoxic environments that trigger glycolytic signaling. Thus, they primarily utilize glycolysis for energy production, whereas mitochondrial oxidative phosphorylation, namely, tricarboxylic acid (TCA) cycle and electron transport chain, is largely compromised. M1 TAMs also upregulate lipid biosynthesis necessary for executing inflammatory responses. On the other hand, M2-TAMs lack a sufficient supply of glucose for glycolysis due to the high glucose consumption by tumor cells. Thus, they resort to mitochondrial oxidative phosphorylation and fatty acid oxidation as the major sources of energy production (Thapa and Lee, 2019).

Interestingly, recent studies have unraveled that these distinct metabolic propensities of M1- vs. M2-TAMs are largely contributed by their differential arginine metabolism. In M1-TAMs that upregulate NOS2, arginine is primarily converted to NO. NO is found to induce S-nitrosylation (conjugation of NO to a cysteine residue) and inhibition of several iron-sulfur proteins present in mitochondrial electron transport chain, such as Complex I and cytochrome c oxidase. This renders M1-TAMs heavily dependent on glycolysis for energy production (Bailey et al., 2019). Conversely, in M2-TAMs that upregulate arginase/ornithine decarboxylase, arginine is preferentially converted to ornithine and then PAs. One of such PAs, spermidine, is found to induce hypusination (conjugation of spermidine to a lysine residue) and activation of eukaryotic initiation factor 5A (eIF5A), the protein responsible for the translation of several enzymes involved in the mitochondrial TCA cycle, such as succinyl-CoA synthetase, succinate dehydrogenase, methylmalonyl-CoA mutase, and pyruvate dehydrogenase. This allows M2-TAMs to resort to mitochondrial oxidative phosphorylation for energy production when there is not enough supply of glucose for glycolysis (Puleston et al., 2019).

Given that distinct arginine metabolites help induce the formation of different TAM types, we hypothesized that a shift of arginine metabolism might trigger TAM reprogramming. In the present study, we tested whether supplementing SEP, the endogenous precursor of the NOS cofactor BH_4_, would redirect arginine metabolism towards NO synthesis and induce reprogramming of M2-TAMs to M1-TAMs. This was based on the previous findings of ours and others that BH_4_ biosynthesis is activated in response to the stimuli involved in M1-TAM polarization (namely, TNFα, IFN γ, and LPS), but is downmodulated in M2-TAMs (Antoniades et al., 2011; Zheng et al., 2020). SEP is a cell permeable pteridine originally discovered as a yellow eye pigment of Drosophila melanogaster (Rao and Cotlier, 1985). SEP is ubiquitously synthesized from bacteria to mammals as part of the salvage pathway of BH_4_ biosynthesis (Nichol et al., 1983). Administering SEP is found to be much more efficient in elevating the intracellular BH_4_ levels than administering BH_4_ itself (Hasegawa et al., 2005; Yamamoto et al., 1996). SEP has been tested as a therapeutic for different metabolic disorders, such as diabetes, hypertension and cardiovascular diseases (Ishikawa et al., 2016; Jones Buie et al., 2019). It has also been utilized for treating phenylketonuria (PKU), the condition defective in phenylalanine catabolism, in phase III clinical trials (Bratkovic et al., 2022; Pannirselvam et al., 2003; Smith et al., 2019; Yoshioka et al., 2015). We recently reported that supplementing SEP could shift arginine metabolism from PA to NO synthesis in both HER2-positive breast cancer cells and TAMs, inhibiting the growth of mammary tumor cells (Ren et al., 2019; Zheng et al., 2020). In the present study, we further explored how SEP induces reprogramming of M2- to M1-TAMs as well as its therapeutic efficacies for promoting T-cell mediated anti-cancer immunity. We showed here that the addition of SEP alone was sufficient to elevate NO to PA ratios in both nascent M0-TAMs and immune-suppressive M2-TAMs and polarize them towards M1-TAMs. These repolarized TAMs manifested bona fide M1-TAM functions, including the full-fledged capabilities of antigen presentation, T effector cell activation, and induction of ICD of HER2-positive tumor cells. We confirmed the pro-immunogenic, anti-tumor effects of SEP using an animal model of spontaneous HER2-positive mammary tumors.

While the present study specifically focuses on the utility of SEP in elevating the immunogenicity of TAMs in HER2-positive breast cancer, the SEP/BH_4_ pathway could also be utilized for the activation or protection of other cell types. For example, the SEP/BH_4_ pathway is critically involved not only in T cell proliferation for anti-tumor activities (Cronin et al., 2018), but also in protection of vascular functions in response to inflammation-induced endothelial damages (Antoniades et al., 2011). Furthermore, because BH_4_ serves as the cofactor of several different enzymes (NOS, phenylalanine/tyrosine/tryptophan hydroxylases, and alkylglycerol monooxygenase), it could exert additional beneficial effects on energy metabolism, redox mechanisms, and disease treatments independent of NOS (Eichwald et al., 2023; Kim and Han, 2020). Besides, both BH_4_ and SEP are FDA-approved drugs (sold commercially under the names sapropterin and PTC923, respectively), and their ‘off-label uses’ have tremendously grown during the last two decades through a number of clinical trials for various conditions including cardiac, pulmonary, rheumatologic, dermic, and psychiatric diseases, dementia, menopause, aging, and inherited disorders (Eichwald et al., 2023). Our study findings also strongly suggest that clinical uses of BH_4_ and SEP for cancer treatment would warrant further investigation.

## Materials & Methods

### Cell lines

Human monocytic cell line THP–1 (Cat. No. TIB-202™), SKBR3 (Cat. No. HTB-30™) and BT- 474 (Cat. No. HTB-20™) cancer cell lines were purchased from American Type Culture Collection (ATCC). CA1d breast cancer cells were obtained from Karmanos Cancer Institute (Detroit, MI) under Material Transfer Agreement. Peripheral blood mononuclear cells (PBMCs) were obtained from Stemcell Technologies (Vancouver, BC, Canada) and AllCells LLC (Alameda, CA).

### Cell culture & Reagents

THP–1 cells were maintained at a density of 1×106 cells/ml in RPMI 1640 Medium (ThermoFisher, Waltham, MA, Cat. No. 11835055) supplemented with 10% fetal bovine serum (FBS), 1% Penicillin/Streptomycin, 2mM GlutaMAX™, 10mM HEPES buffer, 45g/L Glucose and 1mM Sodium Pyruvate (ThermoFisher, Cat. No. 15140122, Cat. No. 35050061, Cat. No. SH3023701, Cat. No. A2494001 & Cat. No. 11-360-070). SKBR3 and BT-474 cells were cultured in McCoy’s 5A Medium with 10% FBS and 1% Penicillin/Streptomycin and CA1d cells were cultured in DMEM/F12 medium with 5% Horse serum, 1% Penicillin/Streptomycin, Hydrocortisone, Cholera Toxin, and Insulin (Sigma-Aldrich, Inc, St. Louis, MO, Cat. No. H-0888, Cat. No. C8052-2MG & Cat. No. I1882). All the cells were maintained in a 37°C humidified incubator with 5% CO2.

### Modulation of Arginine metabolism

For induction of NO, we used NO donors: S-nitroso-N-acetyl penicillamine (SNAP, 10 or 20μM, ThermoFisher, Cat. No. N7892); and S-Nitrosoglutathione (GSNO, 100 or 200μM, Santa Cruz Bio, Dallas, TX, Cat. No. sc-200349B), or SEP, a precursor of NOS cofactor tetrahydrobiopterin: (20 or 100 μM, Career Henan Chemical Co). For NO inhibition, we used Nω-Nitro-L-arginine methyl ester hydrochloride (L–NAME, 2.5 mM, Sigma, Cat. No. N5751-10G); the NOS2 inhibitor: 1400W hydrochloride (100μM, Cayman Chemical, Ann Arbor, MI, Cat. No. 81520); or the NO scavenger: 4-carboxyphenyl-4,4,5,5-tetramethylimidazoline-1-oxyl-3-oxide (cPTIO, 50 or 100 μM, Enzo, Cat. No. ALX-430-001-M050). For induction of PA, we treated cells with Spermine (5 or 10mM, Sigma, Cat. No. 55513-100MG). For inhibition of PA synthesis, we used the polyamine analog, N1,N11-Diethylnorspermine tetrahydrochloride (DENSPM, 50 or 100μM, Tocris, Minneapolis, MN, Cat. No. 0468/10). For inhibition of Arginase 1 (Arg1), we used of N-hydroxy-nor-L-arginine (nor-NOHA, 50 μM, Cayman Chemical, Ann Arbor, MI, Cat. No. 10006861); for inhibition of Ornithine decarboxylase 1 (ODC1), we used Difluoromethylornithine (DFMO, 50 μM, Sigma, Cat. No. D193-25MG).

### In vitro model of TAMs

#### THP–1 derived model

Human monocytes THP–1 cells were seeded at a density of 3×10^5^ cells/ml and treated with 100 ng/ml phorbol myristate acetate (PMA, InvivoGen, San Diego, CA, Cat. No. tlrl-pma) for 24 hours for their differentiation to nascent (M0) macrophages. M0 cells were then serum starved for 2 hours in X-VIVO™ hematopoietic cell medium (Lonza, Basel, Switzerland, Cat. No. BEBP04- 744Q). For M1 polarization (M1-TAMs), M0 cells were treated with PMA (100 ng/ml), 5 ng/ml lipopolysaccharide (LPS, Sigma-Aldrich, Cat. No. L4391-1MG), and 20 ng/ml interferon γ (IFNγ, PeproTech, Cranbury, NJ, Cat. No. 300-02) for 66 hours. For M2 polarization (M2-TAMs), M0 cells were treated with PMA (100 ng/ml), 20 ng/ml interleukin 4 (IL4, PeproTech, Cat. No. 200- 04), and 20 ng/ml interleukin 13 (IL13, PeproTech, Cat. No. 200-13) for 66 hours.

### PBMC derived model

PBMCs were plated at a density of 1.5×10^6^ cells/ml in RPMI serum free media and incubated for 3 hours. Monocytes were isolated based on their adhesion to plastic. Serum free media was aspirated to remove non–adherent cells. The adherent cells were replenished with RPMI media containing 20% FBS and incubated for 24h. Non–adherent cells were further removed by washing with pre–warmed RPMI medium. To induce M0 differentiation, the remaining adherent cells were treated with 50 ng/mL recombinant SCF, and 50 ng/mL granulocyte macrophage colony stimulating factor (GM–CSF, for M1 polarization) or 50 ng/mL macrophage colony-stimulating factor (M–CSF, for M2 polarization) (BioLegend, San Diego, CA, Cat. No. 573904, Cat. No. 572903 and Cat. No. 574804) for 7 days with 50% medium replenishment every 3 days. Upon differentiation, M0 cells were starved for 2 hours as described above and given M1 or M2 polarization treatment as described above for 3 days.

### Reprogramming of M2–TAMs to M1-TAMs

THP–1 and PBMC derived M2–TAMs were treated with 100μM SEP (Career Henan Chemical Co., Zhengzhou City, China, CAS No. 17094-01-8) every day for 3 days for reprogramming to M1 TAMs. M2–TAMs were also treated with DMSO (Vehicle) and 5 ng/ml LPS and 20 ng/ml IFNγ as negative and positive controls, respectively.

### PBMC derived T cell culture

PBMCs were incubated with 100 μg/mL DNase I solution (Stemcell Technologies, Cat. No. 7900) at room temperature (RT) for 15 minutes and filtered through a 37 μm cell strainer. The filtered single cell suspension was centrifuged at 1000 rpm for 5 minutes and the collected pellet was resuspended at a density of 5 x 107 cells/mL in the suspension medium composed of 2% FBS and 1mM ethylenediaminetetraacetic acid (EDTA) in PBS. T cells were then isolated using EasySep™ Human T Cell Isolation Kit and EasySep™ Magnet (Stemcell Technologies, Cat. No. 17951 and Cat. No. 18000) following manufacturer’s protocol. Isolated T cells were seeded at a density of 1 x 106 cells/mL in RPMI medium supplemented with 10 ng/mL human recombinant interleukin 2 (IL2, Stemcell Technologies, Cat. No. 78145.2), 1µg/ml anti-human CD3 antibody (BioLegend, Clone OKT3, Cat. No. 317326) and 1µg/ml anti-human CD28 antibody ((BioLegend, Clone CD28.2, Cat. No. 302934). The cells were maintained in a 37°C, 5% CO2 humidified incubator for up to 3 subcultures with medium change every 3 days.

### Scanning Electron Microscopy (SEM)

THP–1 derived TAM subsets were plated on CellQART® 12-well cell culture inserts with 0.4 µm PET-membrane (STERLITECH, Auburn, WA, Cat. No. 9310412) and treated with SEP and relevant controls. Upon treatment, the membranes were washed with 1X phosphate buffer saline (PBS) and fixed with 4% paraformaldehyde (PFA). Then the membranes were washed with deionized (DI) water and dehydrated using increasing concentrations of ethanol ranging from 25% to 100%. Afterwards, the membranes were washed with 50% Hexamethyldisilazane (HMDS): Ethanol solution followed by 100% HMDS. Later the membranes were removed from the inserts, bound to the SEM sample stub with carbon tape and sputter coated with gold for 15s. The samples were then imaged using JEOL 7500F Scanning Electron Microscope.

### Phalloidin Staining

THP–1 cells were seeded at the density of onto 12 mm coverslips (Thomas Scientific, Swedesboro, NJ, Cat. No. 1217N67), precoated with Poly-D-Lysine (Fisher Scientific, Pittsburgh, PA, Cat. No. A003E) and PureCol® (AdvancedBioMatrix, Carlsbad, CA, Cat. No. 5005-100ML) and singly placed into each well of 12 well plates. These cells were given SEP of control treatment for, washed in 1X PBS and fixed with 3.7% formaldehyde in PBS for 10 minutes at RT. Cells were permeabilized with 0.1% Triton X-100 for 5 minutes and washed twice with PBS. After blocking in PBS containing 1% bovine serum albumin (BSA)for 30 minutes, cells were incubated for 20 minutes at RT with the phalloidin staining solution: Alexa Fluor™ 488 Phalloidin (ThermoFisher, Cat. No. A12379) and 1% BSA in PBS. Cells were washed with PBS and counterstained with 4′,6-diamidino-2-phenylindole (DAPI) nuclear stain for 15 minutes at RT. Coverslips were then mounted on slides using Fluoromount-G Mounting Medium (Fisher Scientific, Cat. No. 50-187- 88) and sealed with nail polish. Fluorescence images were captured using Zeiss Axio Observer 7 microscope.

### Immunofluorescence staining

THP–1 cells were plated on glass coverslips, polarized to TAMs, and given drug treatments as described above. Upon treatment, the coverslips were washed with 1X PBS, and cells were fixed in 4% PFA for 20 minutes at RT. The cells were permeabilized with 0.1% Triton X-100 for 15 minutes at RT and blocked with IF buffer (1.3M NaCl, 70mM Na2HPO4, 35mM NaH2PO4, 77mM NaN3, 1% BSA, 2% Triton X-100 and 5% Tween-20) containing 10% goat serum for 1 hour. Cells were washed with PBS and incubated overnight at 4°C with primary antibody solution: Antibody in IF buffer containing 3% Saponin (Fisher Scientific, Cat. No. 55-825-5100GM). Antibodies used were as follows: TNFα (ThermoFisher, Cat. No. MA523720), NOS2 (ThermoFisher, Cat. No. PA1-036), CD163 (Abcam, Waltham, MA, Cat. No. ab182422), CD206 (Novus Biologicals, Centennial, CO, Cat. No. NPB1-90020), IL10 (R&D systems, Minneapolis, MN, Cat. No. MAB9184-100) and Dectin 1 (ThermoFisher, Cat. No. PA583996). Cells were then washed with PBS twice and incubated with secondary antibody solution (1:200 dil) for 2 hours followed by DAPI stain for 15 minutes at RT. Coverslips were then mounted onto slides as described above. Fluorescence images were captured using Olympus DP80 microscope and analyzed using ImageJ software.

### Immunoblotting

Cell lysates were prepared using the following lysis buffer: 25mM Tris–HCl (pH 8), 150mM NaCl, 1mM EDTA (pH 8), 1% NP–40, 5% Glycerol, 1X PhosSTOP (Sigma-Aldrich, Cat. No. 4906837001) and 1X Protease Inhibitor Cocktail (ThermoFisher, Cat. No. 78425). Total protein concentration of the cell lysates was quantified using the Pierce BCA Protein Assay Kit (ThermoFisher, Cat. No. 23227). Cell lysates were mixed with the sample buffer (with beta mercaptanol) boiled at 100°C for 10 minutes. Proteins were separated by sodium dodecyl-sulfate polyacrylamide gel electrophoresis (SDS-PAGE). The separated proteins were then electroblotted to methanol–activated polyvinylidene fluoride (PVDF) membranes (Sigma-Aldrich, Cat. No. IPVH00010). Upon transfer, membranes were blocked with 5% non-fat milk in Tris-buffered saline containing 0.1% Tween® 20 (TBST) and incubated overnight with the following primary antibodies: TLR2 (ThermoFisher, Cat. No. MA532787), STAT1 (Cell Signaling Technology, Danvers, MA, Cat. No. 9172T), pSTAT1S727 (Cell Signaling Technology, Cat. No. 8826S), CD163 (Abcam, Cat. No. ab182422), CD206 (Novus Biologicals, Cat. No. NPB1-90020), STAT3 (Cell Signaling Technology, Cat. No. 9139S), pSTAT3Y705 (Cell Signaling Technology, Cat. No. 9145S), β–Actin (Sigma-Aldrich, Cat. No. A1978) and GAPDH (Cell Signaling Technology, Cat. No. 2118S). Then they were incubated with horseradish peroxidase (HRP) conjugated Sheep anti-Mouse IgG (GE Healthcare Life Sciences, Pittsburgh, PA, Cat. No. NA931-1ML), Donkey anti-Rabbit IgG (GE Healthcare Life Sciences, Cat. No. NA934-1ML) or Donkey anti-Goat IgG (ThermoFisher, Cat. No. A16005) secondary antibodies (1:5000 dil). Next, the blots were developed with SuperSignal™ West Dura Extended Duration Substrate (ThermoFisher, Cat. No. 34076) and imaged using Syngene G:BOX F3 gel doc system.

### Flow cytometry (FACS) analysis of cell surface markers

Cells were dissociated from the plates through incubation with PBS containing 5mM EDTA for 15 minutes at 37°C, followed by gentle scraping. Cells were collected into 96 well V bottom plates (USA Scientific, Ocala, FL, Cat. No. 5665-1101), and centrifuged at 1000 rpm for 5 minutes. Cell pellets were washed with Fluorescence-activated cell sorting (FACS) buffer (PBS, 2% FBS) and blocked for 30 minutes on ice in the blocking buffer: 2% FBS, 2% goat serum, 2% rabbit serum and 10 µg/mL human Immunoglobulin G (IgG). Cells were then incubated with fluorochrome-labeled antibodies prepared in FACS buffer for 1 hour on ice. The antibodies used are as follows: CD68 (BioLegend, Cat. No. 333821), CD40 (BioLegend, Cat. No. 334305), CD80 (BioLegend, Cat. No. 305205), CD163 (BioLegend, Cat. No. 333609), CD206 (BioLegend, Cat. No. 321109), anti-mouse F4/80 (BioLegend, Cat. No. 123118), anti-mouse CD80 (BioLegend, Cat. No. 104706), anti-mouse CD163 (BioLegend, Cat. No. 155306), anti-mouse NK1.1 (ThermoFisher, Cat. No. 61-5941-80) and anti-mouse NKp46 (R& D systems, Cat. No. FAB2225F-025). Cells were washed twice with FACS buffer and resuspended in Dulbecco’s phosphate-buffered saline (DPBS) containing 2% formaldehyde. Samples were assayed on the BD FACSCanto™ II system, followed by analyses with FlowJo software.

### FACS analysis of intracellular markers

Cells were treated with 1μg/ml Brefeldin A (Fisher Scientific, Cat. No. B7450) for 5 hours at 37°C. Upon treatment, cells were collected into 96 well V bottom plates and centrifuged as described above. The samples were then incubated with 1X Fixation/Permeabilization Buffer (R&D systems, Cat. No. FC007) for 12 minutes at 4°C. Following fixation, samples were centrifuged at 1600 rpm for 5 minutes and washed with the following Permeabilization/Washing buffer: PBS, 2% FBS and 0.1% Triton X-100. Samples were then blocked with blocking buffer containing 0.1% Triton X- 100 for 10 minutes. After blocking, samples were incubated with fluorochrome-labeled antibodies prepared in permeabilization/washing buffer for 45 minutes at 4°C. The following antibodies were used: Anti-mouse IFNγ (R&D systems, Cat. No. IC485F-025), anti-mouse IL12 (BioLegend, Cat. No. 505206), anti-mouse IL10 (BioLegend, Cat. No. 505006) and anti-mouse Perforin (ThermoFisher, Cat. No. 11-9392-80). Following antibody incubation, samples were washed with permeabilization/washing buffer, resuspended in DPBS containing 2% formaldehyde and assayed on the BD FACSCanto™ II system, followed by analyses with FlowJo software.

### Measurement of BH_4_ production

THP–1 derived TAMs plated in 6 wells were used for the measurement of BH4. Cells were washed with ice cold PBS to remove remaining media. Upon washing, cells were scraped gently, and the cell pellets were collected to Eppendorf™ tubes. The cell pellets were vortexed for 10 seconds and flash frozen in liquid nitrogen (N2). Then the pellets were thawed at RT. The process was repeated 5 times. Then the pellets were centrifuged at 1000 rpm for 5 minutes, and the supernatants were collected into fresh tubes. The BH4 level in cell lysate was measured using an enzyme–linked immunosorbent assay (ELISA) Kit (Abbexa, Sugar Land, TX, Cat. No. abx354211) following manufacturer’s protocol. Cellular BH4 levels were normalized using the total protein concentration in cell lysates.

### Live cell imaging of NO and PA

Intracellular NO and PA levels were detected in live cells with the help of cell permeable fluorescent dyes. Cells grown in phenol red, and serum free media were stained with 20μM Diaminorhodamine-4M acetoxymethyl ester (DAR-4M AM) (Enzo, Farmingdale, NY, Cat. No. ALX-620-069-M001) or 20μM PolyamineRED (Diagnocine, Hackensack, NJ, Cat. No. FDV- 0020) according to the manufacturers’ protocol to detect NO and PA respectively. Fluorescence images were captured using Olympus DP80 microscope and analyzed using ImageJ software.

### Measurement of NO levels

Nitric Oxide Fluorometric Assay Kit (BioVision, Catalog no. K252) was used to quantify the NO levels present in the CM. The samples were filtered through a 10 kDa cut-off Microcon filter (Millipore, St. Louis, MO, USA) to remove proteins present in the CM. The flow through was reacted with the assay reagents in the dark according to the manufacturer’s protocol and the fluorescence intensity was measured at the wavelengths of Ex/Em = 360/450 nm.

### Measurement of PA levels

PA levels in the CM were quantified using Fluorometric Total Polyamine Assay Kit (BioVision, Catalog no. K475) according to the manufacturer’s protocol with modifications. This kit determines the level of hydrogen peroxide produced through oxidation of polyamines by spermine/spermidine oxidase in the kit. To remove high background levels of hydrogen peroxide produced by macrophages prior to the assay, the CM were pretreated with Catalase (Sigma-Aldrich, Cat. No. C40-100MG) at 100 μg/ml and incubated at 37°C for 1.5 hrs. Proteins were precipitated with Sample Clean-up Solution provided by the kit and removed by filtration through a 10 kDa cut-off Microcon filter. The flow through was reacted with the assay reagents in the dark according to the manufacturer’s protocol and the fluorescence intensity was measured at the wavelengths of Ex/Em = 535/587 nm.

### Measurement of cytokine secretion

Cells were cultured in RPMI serum free media for 2 days following treatment and the CM were collected. Secreted cytokines in the CM were quantified using ELISA kits according to manufacturers’ protocol. The ELISA kits used were as follows: IL12 (R&D systems, Cat. No. D1200), IL6 (R&D system, Cat. No. D6050), IL1β (R&D system, Cat. No. DLB50), TNFα (R&D system, Cat. No. DTA00D), IL10 (R&D systems, Cat. No. D1000B) and TGFβ (Abcam, Cat. No. ab108912).

### Antigen presentation assay

THP–1 derived TAMs were detached from the plates, collected into 96 well V bottom plates and centrifuged as described before. The cells were then pulsed with 20μg/ml OVA323–339 peptide (GenScript, Piscataway, NJ, Cat. No. RP10610-1) for 2 hours at 37℃ and 5% CO2. Cells were then washed twice with FACS buffer (see above) to remove residual OVA peptides and incubated with blocking buffer for 30 minutes at 4°C. Samples were stained with fluorescein isothiocyanate (FITC) labeled HLA–DR antibody (Biolegend, Cat. No. 307604) in FACS buffer for 1 hour at 4°C and analyzed by flow cytometry.

### TAM, T cell and cancer cell co–culture model

A triple co–culture model of breast cancer cells, TAMs and T cells was developed to emulate and study the tumor–immune interactions within the TME.

### Transwell co–culture system

THP–1 cells were seeded on CellQART® 12-well inserts (0.4 µm pore size and PET-membrane) at a density of 3×105 cells/ml and subjected to differentiation, polarization, and reprogramming as described above. After establishment of macrophages, the inserts were washed with fresh RPMI media, and breast cancer cells (CA1d, SKBR3 or BT-474) in RPMI media were seeded on top of THP-1 cells at a ratio of 1:2 (cancer cells: THP-1 cells). Into separate bottom wells of 12 well plates, the equal amounts of breast cancer cells were seeded in RPMI media. Once cells were attached, PBMC derived T cells were seeded on top of cancer cells at a ratio of 1:2 (cancer cell: T cell). The inserts containing cancer cells and THP-1 were placed on top of the wells containing cancer cells and T cells. The co–cultures were incubated in a humidified incubator set at 37°C with 5% CO2 for further experiments.

### Direct co–culture system

Monocytes and T cells isolated from the same PBMCs were used to establish the direct co–culture system. Monocytes were seeded on 12 well plates at a density of 1.5×106 cells/ml, differentiated, polarized into relevant TAM subsets, and reprogramed as described above. The media were removed, and cells were washed with fresh RPMI serum free media. Breast cancer cells and T cells were sequentially added to the 12 well plates at a ratio of 1:2:2 (cancer cell: TAM: T cell) in the same RPMI media. The co–culture was incubated in a humidified incubator set at 37°C with 5% CO2 for further experiments.

### T cell proliferation assay

PBMC derived T cells were washed with PBS and resuspended at a density of 1x106 cells/ml in. The cells were then stained with the CellTrace™ CFSE dye (ThermoFisher, Cat. No. C34554) following manufacturer’s protocol. Carboxyfluorescein succinimidyl ester (CFSE) labeled T cells were directly co–cultured with PBMC derived TAMs and cancer cells as described above and incubated for 4 days. T cells were isolated and stained with fluorochrome-labeled anti human CD3 (BioLegend, Cat. No. 317342) and anti human CD8a (BioLegend, Cat. No. 301014) antibodies as described above. The cytotoxic T cell proliferation, indicated by changes in CFSE signals, was measured by flow cytometry.

### Measurement of IFNγ production

The triple co–culture, composed of cancer cells, TAMs and T cells, was established as described above and incubated for 2 days. Following incubation, the cells were treated with 1μg/ml Brefeldin A for 4 hours to block protein secretion. T cells were isolated and stained with anti-human CD3 and CD8a antibodies (surface staining) followed by intracellular staining with anti- human IFNγ (R&D Systems, Cat. No. IC285F-100) antibody, as described before, and analyzed by flow cytometry.

### Degranulation assay

CD107a marker expression was measured to analyze the degranulation of cytotoxic T cells [1, 2]. The triple co–culture system was set up as previously described and incubated for 2 days. Upon incubation, 10μl of anti-human CD107a antibody (BioLegend, Cat. No. 328606) was added to each well and incubated in a humidified incubator set at 37°C with 5% CO2. After 1 hour of incubation, 1X eBioscience™ Monensin Solution (Fisher Scientific, Cat. No. 00-4505-51) was added to each well and the incubation for 3 more hours to block protein transport. T cells were then isolated and stained with CD3 and CD8a antibodies, as previously described. CD107a expression in cytotoxic T cells was measured by flow cytometry.

### Measurement of extracellular ATP

During immunogenic cell death (ICD) — a form of apoptosis, dying cells release ATP to induce immunogenic phagocytosis. Thus, the increase in secreted ATP is used to determine the occurrence of ICD [3, 4]. Luminescent ATP detection assay kit (Abcam, Cat. No. ab113849) was used to quantify ATP released from cells. The co–culture system was set up as described above, and 100μl of CM from the bottom well (seeded with cancer cells and T cells) was analyzed for ATP levels, in comparison to CM of monocultured TAMs, T cells and cancer cells as controls. ATP concentration was quantified using the standard following manufacturer’s protocol.

### Cell Cycle Analysis

Cancer cells were labeled with CellTracker™ Green CMFDA Dye (ThermoFisher, Cat. No. C7025) according to manufacturer’s protocol. The labeled cancer cells were directly co–cultured with PBMC-derived TAMs and T cells as described above and incubated for 4 days. Cells were harvested using 0.05% Trypsin (Fisher Scientific, Cat. No. 25-300-062), washed with PBS, and centrifuged at 1000 rpm for 5 minutes. The pelleted cells were fixed with pre–cooled 70% ethanol for 2 hours at 4°C. Cells were centrifuged at 4000 rpm for 2 minutes, resuspended in PBS containing 0.25% Triton X–100, and incubated on ice for 15 minutes. Cells were centrifuged and resuspended in PBS containing 10 μg/ml Ribonuclease (RNase) A and 20 μg/ml Propidium Iodide (PI) (Sigma, Cat. No. R4642-10MG and Cat. No. P4170-10MG) for the staining of DNA. Cells were transferred to FACS tubes, incubated in the dark for 30 minutes at RT, and analyzed for cell cycle profiles by flow cytometry.

### Apoptosis Assay

The direct co–culture model was set up using CMFDA labeled cancer cells as described before and incubated for 4 days. Adherent cells were harvested, washed with FACS buffer, and centrifuged at 1000 rpm for 5 minutes. The pelleted cells were then resuspended at a density of 1x106 cells/ml in Annexin V Binding Buffer: 10mM HEPES, 150mM NaCl and 2.5mM CaCl2 in PBS. Resuspended cells were transferred to FACS tubes, stained with 5μl of Annexin V (ThermoFisher, Cat. No. A35110) and 20 μg/ml of PI and incubated in the dark for 15 minutes at RT. Each tube was replenished with 400 µL of Annexin V Binding Buffer and analyzed by flowcytometry.

### Animal Study

All in vivo experiments were performed in compliance to The Guide for the Care and Use of Laboratory Animals (National Research Council, National Academy Press, Washington, D.C., 2010) and with the approval of the Institutional Animal Care and Use Committee of the University of Toledo, Toledo, OH (Protocol No: 108658). Two months old female MMTV-neu (unactivated) (n=14) mice were obtained from the Jackson Laboratory (ID. IMSR_JAX:002376, Bar Harhor, MN, USA), housed under regular conditions and given ad libitum access to acidified water and regular chow. The mice were maintained until spontaneous mammary tumors became palpable (∼ 5 mm long, 6-14 months). Upon tumor onset, mice were divided into vehicle (DMSO) vs. SEP (10mg/kg) treatment groups (Pannirselvam et al., 2003; Yoshioka et al., 2015). The drugs were dissolved in acidified drinking water and administered to mice ad libitum for 6 weeks (Rabender et al., 2015). The tumor growth, body weight and the morbidity of the animals were monitored twice a week. At the end of treatment, mice were euthanized, and mammary tumors were processed for further analyses described below.

### Profiling of macrophages cells in tumors and spleens

Freshly harvested tumors and spleens were processed for the profiling of resident macrophages. Tumors and spleens were weighed and reacted in the digestion mixture [10ml/g of tissue, 3mg/ml Collagenase A (Sigma, Cat. No. 10103578001) and 25μg/ml DNase I (Sigma, Cat. No. 10104159001)] in Hanks’ Balanced Salt Solution (HBSS, ThermoFisher, Cat. No. 14025092) with gentle motions on a platform shaker for 45 minutes at 37°C (Krneta et al., 2016). To stop the enzymatic digestion, the samples were treated with 10 ml of staining buffer (1% (w/v) BSA in PBS). Cell suspension was then filtered through a 100 μm cell strainer, and the volume was adjusted to 20 ml with staining buffer. Then, cells were pelleted by centrifugation at 500g for 7 minutes at 4°C. To remove red blood cells, the pelleted cells were suspended in 3 ml of 1X Red Blood Cell (RBC) lysis buffer (ThermoFisher, Cat. No. 00-4300-54) and incubated on ice for 10 minutes. A volume of 30 ml of staining buffer was added, cells were centrifuged and resuspended in 1ml of staining buffer. To block Fc receptors (to avoid unwanted antibody binding), cells were treated with 2 μl of anti-mouse CD16 (FcγII)/CD32 (FcγIII) Antibody (ThermoFisher, Cat. No. 14-0161-82) and incubated on ice for 30 minutes with mixing at 10 minute intervals. A volume of 4 ml of staining buffer was added, and samples were centrifuged at 500g for 5 minutes at 4°C. Then, cell pellets were resuspended in 1ml of staining buffer (Cassetta et al.). Cells were then stained with relevant fluorochrome labeled antibodies, as previously described, and analyzed by flow cytometry.

### Statistical Data Analysis

The experimental results are presented as mean ± SEM. All the experiments were performed in replicates (n ≥ 3) and, unless otherwise indicated, two-tailed t-tests were performed to obtain the statistical significance of the mean difference. P values ≤ 0.05 were considered statistically significant. Flow cytometry data analyses were performed using FlowJo Version 10.5. Imaging analyses were performed on ImageJ software. All statistical analyses were carried out using GraphPad Prism Version 9.5.1.

## Acknowledgement

We thank Dr. Eun-Seok Choi in Furuta laboratory at Case Western Reserve University for manuscript review and constructive suggestions. We would also like to thank Drs. Andrea Kalinoski and David Weaver in the Imaging Core at the University of Toledo for various supports in imaging and FACS analyses.

## Author Contributions

Conceptualization, V.F and S.F; Methodology, V.F, X.Z., V.S. and S.F; Formal Analysis: V.F, X.Z., and V.S; Investigation, V.F, X.Z., and V.S.; Data Curation, V.F, X.Z., and V.S; Writing – Original Draft Preparation, V.F. and S.F.; Visualization, V.F, V.S and S.F.; Supervision, S.F.; Project Administration, S.F.; Funding Acquisition, S.F.

## Supporting Information

**Appendix (PDF document)**

